# Swine viral detection by adapted Next-Generation Sequencing (NGS) for RNA and DNA species reveals first detection of porcine circovirus type 3 (PCV3) in Chile

**DOI:** 10.1101/2020.06.07.138925

**Authors:** Paulina S. Rubilar, Javier Tognarelli, Jorge Fernández, Cristóbal Valdés, Felipe Broitman, Dinka Mandakovic, Rodrigo Pulgar

**Author notes:** Corresponding author: Rodrigo Pulgar, Laboratorio de Genómica y Genética de Interacciones Biológicas (LG2IB), INTA, Universidad de Chile, Santiago, Chile.

## Abstract

The outbreak and propagation of COVID-19 have posed a significant challenge to our society, highlighting the relevance of epidemiological monitoring of animal pathogens closely related to humans due to potential zoonosis. To find previously undetected viruses inside Chilean swine farms, we collected non-invasive samples from animals suspected to undergo any viral disease. After screening by Next Generation Sequencing (NGS) adapted to identify viral species with RNA and DNA genomes simultaneously, different viral species were successfully detected. Among viruses with an RNA genome, Porcine Rotavirus A (RVA), Porcine Astrovirus type 5 (PAstV-5) and Porcine Feces-Associated IASV-like virus (PfaIV) were identified; whereas among viruses with DNA genome, Porcine Parvovirus 1 (PPV1), Porcine Circovirus type 1 (PCV1) and Porcine Circovirus type 3 (PCV3) were found. PCV3, a recently described pathogenic virus, was detected for the first time in Chile in different samples from mummified tissue of stillbirths. The whole genome sequence of PCV3 was completed using conventional PCR and Sanger sequencing and compared with previously reported genomes. To the best of our knowledge, this is the first time that an adapted NGS system is used for the screening of DNA and RNA viral species simultaneously inside the Chilean swine industry. This adapted NGS system might be useful for the detection of swine viral emerging diseases worldwide.

---

The outbreak and propagation of COVID-19 have posed a significant challenge to our society, highlighting the relevance of epidemiological monitoring of animal pathogens closely related to humans due to potential zoonosis^1^. Molecular diagnostics have proven to be useful for accelerating detection of ongoing diseases both in humans and in animals^2^. Diagnostics and surveillance of human pathogens in developed countries is mostly entrusted to protocols involving NGS in order to face diseases faster and avoid epidemics. Likewise, international animal diagnostics using NGS are growing, especially for detection of emergent viral pathogens^3,4^. This has been the case for swine re-emerging pathogens causing the African Swine Fever or the Porcine Reproductive and Respiratory Syndrome, viruses that recently have caused severe economic losses worldwide^5^. Chile possesses natural geographic borders with neighboring countries that limit animal displacement. However, the Chilean swine industry is vulnerable to epidemics due to the import of animals for breeding. Progenitors are usually obtained from countries of the north hemisphere, which constitute a probable route for the introduction of emerging pathogens despite quarantine of all animals introduced to the country. Viral infectious diseases are often neglected unless outbreaks emerge, and screening of international pathogens during or after quarantine is not currently performed. On the one hand, most reports on animal viral detection by NGS to date are standardized for single virus detection. In simplified terms, these approaches usually comprise extraction of either DNA or RNA viral genomes followed by specific amplification of viral copies with specific oligonucleotides before or concurrently to the library preparation. Resulting reads are assembled against reference genomes available; thus, less bioinformatic capacity is needed.

On the other hand, sequencing of unknown viral species is a more complex task as viruses differ broadly in genome size, structure, and composition. Besides, more sophisticated algorithms are required for data analysis, such as the assembly of reads “de novo” without reference genomes. A previous study in pigs has shown simultaneous detection of DNA and RNA viruses in feces^6^. The study reported the extraction of total nucleic acids of concentrated viral particles using pooled samples of feces of five healthy pigs and a CsCl density gradient. Amplification of total RNA was used to increase viral copies before NGS, and three types of phages were used as controls. However, ultracentrifuges and complicated methods for viral concentration are usually not a choice in small labs. Thus, before sample processing, it is strongly recommended first that researchers explore equipment accessibility and available samples to compare for time and cost-efficient viral nucleic acids extractions^6,7^. Second, considering that different pipelines for analysis may lead to result discrepancies, it is relevant to choose an adequate platform for the study^6,8^.

In the context of public-private funding search, the use of NGS technology inside the Chilean swine industry was implemented to detect viral species that could be affecting productivity. In the present study, a mix of 30 random octamers previously reported was used to reverse transcribe viral sequences avoiding ribosomal RNA sequences from bacteria and human^9^. A total of 85 samples were collected from 60 animals suspected of undergoing any viral disease (Supplementary table 1). Among clinical signs, there were pigs with diarrhea, flu-like signs, ear-and-tail necrosis, and mummified stillborn fetuses. Most of the samples were non-invasive, including nasal or fecal swabs, semen, placenta or blood. Exceptionally, tissue samples were taken from mummified stillborn fetuses and sick animals of unknown etiology, which were euthanized for merciful reasons. No animal experimentation was made for this study. Samples were transported in DNA/RNA Shield (Zymo Research, Irvine, Ca, USA) at room temperature and stored at −20 °C in the same medium until total nucleic acid extraction. Total nucleic acids (RNA and DNA) were obtained from the samples described above using MagMAX™ CORE Nucleic Acid Purification Kit (Applied Biosystems, Foster City, CA, USA). For NGS, instead of using RNA only as previously described^9^, total nucleic acids were reverse transcribed using Superscript III First-Strand Synthesis Supermix (Invitrogen, Carlsbad, Ca, USA), combined with the mixed 30 octamers reported previously20 at 1μM final each^9^. Second strand DNAs were synthesized using Klenow fragment of DNA PolI (Promega, Madison, WI, USA) followed by isothermal amplification using either Phi29 DNA Polymerase or EquiPhi29™ DNA Polymerase (Thermo Scientific, Waltham, MA, USA) at 30 or 42 °C respectively for 3 hours. Obtained double-stranded DNAs were purified using 2 Volumes of Agentcourt® AMPure® XP (Beckman Coulter, Brea, CA, USA) and followed by quantification using Quant-IT™ DNA Assay Kit, Broad Range (Invitrogen™) and VICTOR® Nivo™ microplate reader (Perkin Elmer, Waltham, MA, USA). Despite all 85 samples undergoing this processing into double-stranded DNA/cDNA, only 70 samples did meet the minimum concentration for DNA library synthesis. Samples were normalized into 2nM concentration, and libraries were prepared using Nextera XT Library Prep Kit (Illumina, San Diego, CA, USA) according to manufacturing instructions. After library purification, samples were pooled and loaded into the MiSeq System (Illumina). For data analysis, raw reads were assessed for quality using FastQC (bioinformatics.babraham.ac.uk/projects/fastqc/). Trimming, to remove adapters and low-quality bases (Q < 10), was performed using BBDuk software (sourceforge.net/projects/bbmap/). De Novo assembly was performed using SPAdes (cab.spbu.ru/software/spades/) or MetaSPAdes using virMine tool, which is useful for the assembly of viral species^10^. When comparing virMine output FASTA files generated for every sample, it was noticed that occasionally, a genuine viral contig found within the “final_contigs” file was missing within the “viral_contigs” file; thereafter all final_contigs files were analyzed for every sample using nucleotide Blast from NCBI. Default parameters or virus txid: 10239 restricted searches of Blast provided similar results considering identity parameters over 90% and query coverages over 60% for every contig.

Careful revision needed to be made for some endoretroviral sequences with identity scores over 90% that also matched host genome sequences with slightly higher identity. The results indicated that viral species were detected in 12 out of the 70 samples (17%). Viral sequences constituted less than 1% from total contigs that matched mainly host sequences or bacteria, depending on the sample type. Among positive samples, 3 (25%) contained contigs matching two different viral species. RNA genome viruses were identified. Namely, a fecal swab from an animal undergoing diarrhea matched contigs for Porcine Astrovirus type 5 (PAstV-5) and Rotavirus A (RVA); whereas, the second sample of fecal swab matched contigs for PAstV-5 and Porcine Feces-Associated IASV-like virus (PfaIV). PAstV-5, detected in this study on fecal samples of animals undergoing diarrhea, is a positive-sense single-stranded RNA virus that has been previously reported in feces of both healthy and ill animals^11^. A previous study has already reported simultaneous detection of PAstV and Rotavirus species on diarrheic swine^12^, which may suggest that coinfection of these viruses could be required for causing diarrhea. Additionally, Porcine astroviruses have been reported as neurovirulent, associated with encephalomyelitis outbreaks in different countries^13^ Rotaviruses, on the other hand, are a group of double-stranded RNA viruses with a segmented genome that cause acute gastroenteritis in several species. Porcine rotaviruses have demonstrated to cause several economic losses in the swine industry, with RVA having the highest prevalence^14^; yet, high viral diversity of this family has led to a limited coverage of vaccines and consequently, to reduced immunization practices^15^. Lastly, PfaIV was detected on a feces sample from a 28 days old piglet undergoing severe diarrhea. This virus has been previously detected on pig feces with no direct association to disease, as it was also found in asymptomatic piglet feces^16^.

DNA genome viruses were also identified. Among these viral species, Porcine Parvovirus type 1 (PPV1) was found on a swab sample from the abdominal cavity taken from a grown animal with signs of polyserositis. PPVs are small, non-enveloped, and single-stranded DNA viruses belonging to family Parvoviridae^17^. PPV1 causes reproductive failure, also known as SMEDI syndrome (stillbirths, mummification, embryonic death, and infertility), and does not cause any disease sign in non-pregnant pigs^18^. Also, the coinfection of PPVs and Porcine Circoviruses has been described, suggesting a possible association between these viruses^18^. Moreover, two species of porcine circovirus, also DNA viruses, were detected among Chilean samples. Circoviruses are non-enveloped viruses with a single-stranded circular DNA genome^19^. There are three species of porcine circovirus (PCV1, PCV2, and PCV3), although only PCV2 and PCV3 are considered pathogenic^18^. In this study, we detected contigs matching PCV1 and PCV3 in samples associated with reproductive failure. Coinfection was also detected in a sample of the placenta from a sow delivering mummified stillbirth fetuses. To compare the pathogenic PCV3 identified in this study with previously reported isolates, we completed the sequence of the whole genome (2000 nucleotides). To accomplish this, NGS resulting contigs matching PCV3 obtained from a mummified tissue sample, were aligned with a couple of full genomes obtained from NCBI database. Alignment resulted in approximately 80% of the full genome sequence covered, lacking a middle portion of approximately 400 nucleotides. This remaining 20% was completed using oligonucleotides reported previously, followed by Sanger-sequencing^20^. Phylogenetic and molecular evolutionary analyses were conducted using MEGA6 (www.megasoftware.net) by aligning the full PCV3 genome obtained in Chile with the 318 full genome sequences from 17 different countries (available on NCBI nucleotide database by December 2019). PCV3 genome described in this study is the southernmost genome reported to date (Figure 1A), with close similarity to PCV3b genomes obtained from samples from Italy, South Korea, China, Hungary, and Brazil (Figure 1B). The full genome sequence was submitted to the NCBI repository under the GenBank accession number MN907812. Remarkably, PCV3 was the most frequently detected virus among our samples, even though it has not been previously described in Chile. To confirm this finding, we randomly selected 50 samples among the 85 total samples, including a couple of those positive to PCV3 by NGS as controls and those samples that did not meet the minimum concentration for NGS library synthesis, and tested the amplification by PCR of a 328 bp viral segment of the cap ORF using specific primers reported previously^21^. Specific amplification was confirmed by Sanger sequencing using Big Dye Terminator Kit v3.1 (Applied Biosystems™) and a 3500 Genetic Analyzer (Applied Biosystems, Foster City, CA, USA), followed by nucleotide Blast. The results confirmed the presence of PCV3 in approximately 70% of the samples tested. These samples represent mostly symptomatic animals from 3 different sites geographically separated along the country and belonging to the same company; thus, this high circulating percentage does not apply to all Chilean pig farms. As a result of this finding, further studies will be undertaken across the country with different companies to assess the relevance of PCV3 within the Chilean swine industry. Also, a full genome diversity evaluation for the related pathogen PCV2 will be perfomed, with special focus on Cap protein against which vaccines are currently being developed.

**Figure 1.**
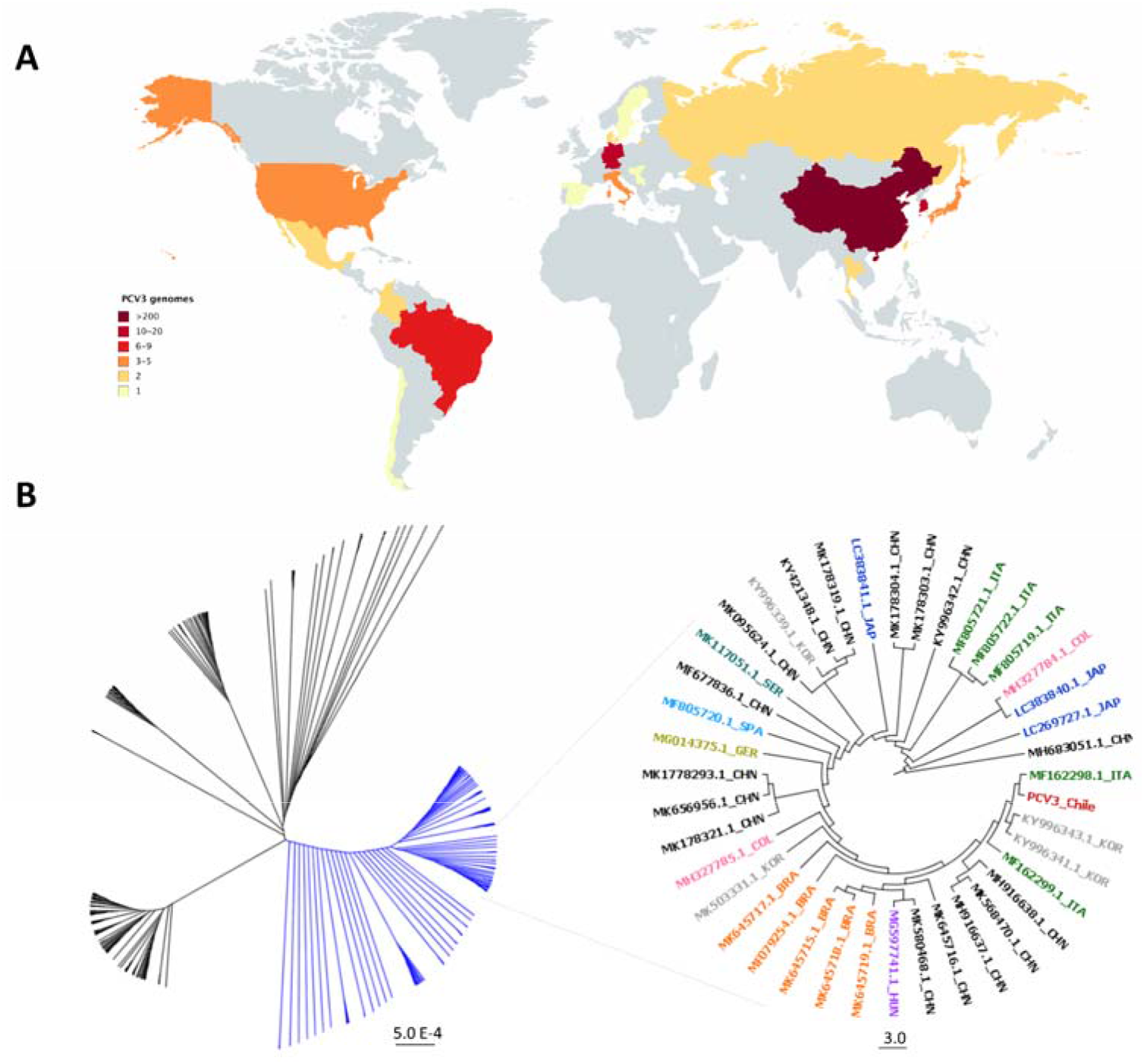
Worldwide distribution of countries and phylogenetic analysis of PCV3 genomes. **A.** Frequency and distribution of countries that have sequenced PCV3 genomes. **B.** Left, UPGMA tree resulting from the analysis of 318 PCV3 genomes sequenced worldwide. Black branches correspond to PCV3a group; blue branches correspond to PCV3b group. Right zoom to clade from PCV3b group that includes the Chilean strain (PCV3_Chile). ITA: Italian strains; KOR: South Korean strains; CHN: Chinese strains; HUN: Hungary strain; BRA: Brazilian strains; COL: Colombian strains; GER: German strain; SPA: Spanish strain; SER: Serbian strain; JAP: Japanese strains. Different colors represent different countries.

Altogether, these results indicate that the adapted procedure for swine viral species reported herein has shown to be useful for the identification of viral pathogens containing either RNA or DNA genomes. Total nucleic acid extraction seemed suitable in this context; however, it contributed to a high presence of host genome sequences that might lead to a lower sensitivity of the approach. These results allowed us to confirm, for the first time, the presence of PCV3 circulating in Chilean swine farms.

Early identification of pathogens is crucial for the decision-making process regarding the prevention, control, and treatment of disease as well as for mitigation of the economic impact of viral diseases. Thus, the availability of more straightforward protocols like the one described in this work using NGS for searching and studying swine viruses is relevant to enhance productivity and to provide the required preparedness to face possible viral outbreaks inside the swine industry.

## Acknowledgments

We would like to thank Daniela Guiñez, who collected most of the samples included in this study, and Vivian Gomez, who processed some Sanger-seq samples.

