## Supplementary table 1 for "Swine viral detection by adapted Next-Generation Sequencing (NGS) for RNA and DNA species reveals first detection of porcine circovirus type 3 (PCV3) in Chile"

**Supplementary table 1**: Description of samples analyzed in this study

ND=not done

| *Sample number* | animal | Sample | Signs | PCR | NGS total contigs | NGS viral |
|---|---|---|---|---|---|---|
| *1* | 1 | Lung tissue | Chess-board lob. Consol., dead | PCV3+ | 210 | none identified |
| *2* | 2 | Large bowel tissue | Diarrhea, dead | PCV3+ | 198 | none identified |
| *3* | 3 | Fetal muscle | Stillborn 1 | PCV3+ | ND | ND |
| *4* | 4 | Fetal muscle | Stillborn 2 | PCV3+ | ND | ND |
| *5* | 5 | Fetal muscle | Stillborn 3 | PCV3+ | 228 | PCV3 (2contigs) |
| *6* | 6 | Fetal muscle | Stillborn 4 | PCV3+ | 203 | PCV3 (4 contigs) |
| *7* | 7 | Serum | Mother of stillborn 1 | PCV3+ | 136 | none identified |
| *8* | 8 | Serum | Mother of stillborn 2 | PCV3+ | ND | ND |
| *9* | 9 | Serum | Mother of stillborn 3 | PCV3+ | 179 | none identified |
| *10* | 10 | Feces | Diarrhea | RVA+ PAstV5+ IASV+ | 789 | RVA (2 contigs) PAstV5(7contigs) |
| *11* | 11 | Feces | Diarrhea | RVA+ IASV+ | 256 | RVA (1 contigs) |
| *12* | 12 | Nasal swab | Coughing | PCV3+ | 163 | none identified |
| *13* | 13 | Nasal swab | Coughing | PCV3+ | 125 | none identified |
| *14* | 14 | Nasal swab | Coughing | PCV3+ | ND | ND |
| *15* | 15 | Nasal swab | Coughing | PCV3+ | 166 | none identified |
| *16* | 16 | Nasal swab | Ear necrosis | PCV3+ | 136 | none identified |
| *17* | 17 | Feces | Diarrhea | PCV3+ IASV+ | 198 | none identified |
| *18* | 18 | Myocardial tissue | Tail and ear necrosis | PCV3+ | 296 | PCV3 (1 contig) |
| *19* | 18 | Auricle tissue | Tail and ear necrosis | PCV3+ | ND | ND |
| *20* | 18 | Spleen | Tail and ear necrosis | PCV3+ | 133 | none identified |
| *21* | 18 | Serum | Tail and ear necrosis | PCV3+ | ND | ND |
| *22* | 19 | Serum | Ear necrosis | PCV3+ | ND | ND |
| *23* | 20 | Serum | Necrosis de cola | PCV3- | ND | ND |
| *24* | 21 | Semen | Healthy male A4799 | PCV3+ | ND | ND |
| *25* | 22 | Semen | Healthy male A4828 | PCV3+ | ND | ND |
| *26* | 23 | Semen | Healthy male A4779 | PCV3+ | ND | ND |
| *27* | 24 | Nasal swab | Flu-like symptoms | PCV3+ | 164 | none identified |
| *28* | 24 | Serum | Flu-like symptoms | PCV3+ | ND | ND |
| *29* | 25 | Nasal swab | Healthy | PCV3+ | 125 | none identified |
| *30* | 25 | Serum | Healthy | PCV3+ | ND | ND |
| *31* | 26 | Nasal swab | Flu-like symptoms | PCV3+ | 153 | none identified |
| *32* | 26 | Serum | Flu-like symptoms | PCV3+ | ND | ND |
| *33* | 27 | Nasal swab | Flu-like symptoms | PCV3- | ND | ND |
| *34* | 27 | Serum | Flu-like symptoms | PCV3+ | ND | ND |
| *35* | 28 | Nasal swab | Flu-like symptoms | PCV3+ | 173 | none identified |
| *36* | 29 | Serum | Healthy | PCV3- | ND | ND |
| *37* | 30 | Feces | Diarrhea | IASV+ | 328 | PAstV5 (2contigs) |

| | | | | | | |
|---|---|---|---|---|---|---|
| | | | | PCV3- | | |
| 38 | 31 | Feces | Diarrhea | IASV+ PCV3- | 248 | PAstV5 (3 contigs) IASV (1contigs) |
| 39 | 32 | Feces | Diarrhea | IASV+ PCV3- | 344 | none identified |
| 40 | 33 | Feces | Diarrhea | IASV+ PCV3- | 429 | PAstV5 (2contigs) |
| 41 | 34 | Lung tissue | Pneumonia | ND | 226 | none identified |
| 42 | 35 | Lung tissue | Pneumonia | ND | 107 | none identified |
| 43 | 36 | Large bowel tissue | Pneumonia | ND | 141 | none identified |
| 44 | 37 | Large bowel tissue | Bowel inflammation | ND | 6 | none identified |
| 45 | 38 | Peritoneum swab | Polyserositis | ND | 168 | PCV1 (2 contigs) |
| 46 | 38 | Lung tissue | Polyserositis | ND | 172 | none identified |
| 47 | 39 | Tracheal swab | Pneumonia, bowel inflammation. | ND | 150 | none identified |
| 48 | 39 | Lung tissue | Pneumonia, bowel inflammation | ND | 172 | none identified |
| 49 | 40 | Semen | Healthy maled13 | ND | 1966 | none identified |
| 50 | 41 | Semen | Healthy male B70 | ND | 1151 | none identified |
| 51 | 42 | Semen | Healthy male B66 | ND | 1018 | none identified |
| 52 | 43 | Semen | Healthy male E10 | ND | 790 | none identified |
| 53 | 44 | Serum | Mother Z811 | ND | 167 | none identified |
| 54 | 44 | Placenta | Mother Z811 | ND | 306 | PAstV5 (2contigs) |
| 55 | 45 | Fetal heart | Stillborn 1 from Z811 | ND | 126 | none identified |
| 56 | 45 | Fetal lung tissue | Stillborn 1 from Z811 | ND | 278 | none identified |
| 57 | 45 | Fetal liver tissue | Stillborn 1 from Z811 | ND | 62 | none identified |
| 58 | 46 | Fetal heart | Stillborn 2 from Z811 | ND | 136 | none identified |
| 59 | 46 | Fetal lung tissue | Stillborn 2 from Z811 | ND | 10 | none identified |
| 60 | 46 | Fetal liver tissue | Stillborn 2 from Z811 | ND | 27 | none identified |
| 61 | 47 | Fetal heart | Stillborn 3 from Z811 | ND | 120 | none identified |
| 62 | 47 | Fetal lung tissue | Stillborn 3 from Z811 | ND | 230 | none identified |
| 63 | 47 | Fetal liver tissue | Stillborn 3 from Z811 | ND | 220 | none identified |
| 64 | 48 | LCR | Polyserositis | ND | 300 | none identified |
| 65 | 49 | Fetal lung tissue | Pneumonia, kidney inflammation. | ND | 223 | none identified |
| 66 | 49 | Thorax exudate swab | Pneumonia, kidney inflammation | ND | 175 | none identified |
| 67 | 50 | Serum | Healthy mother TN1322 (only healthy piglets) | ND | 130 | none identified |
| 68 | 51 | Serum | Healthy newborn A from TN1322 | ND | 205 | none identified |
| 69 | 52 | Serum | Healthy newborn B from TN1322 | ND | 265 | none identified |
| 70 | 53 | Serum | Healthy newborn C | ND | 283 | none identified |

| | | | | | | |
|---|---|---|---|---|---|---|
| | | | from TN1322 | | | |
| 71 | 54 | Serum | Healthy newborn D from TN1322 | ND | 244 | none identified |
| 72 | 55 | Serum | Mother X | PCV3+ | 244 | none identified |
| 73 | 55 | Placenta | Mother X | ND | 241 | none identified |
| 74 | 56 | Fetal heart | Stillborn from X | ND | 472 | none identified |
| 75 | 57 | Serum | Mother TN1960 | PCV3+ | 202 | none identified |
| 76 | 57 | Placenta | Mother TN1960 | PCV3+ | 609 | none identified |
| 77 | 58 | Fetal muscle | Stillborn from TN1960 | PCV3+ | 905 | none identified |
| 78 | 58 | Fetal heart | Stillborn from TN1960 | PCV3+ | 427 | none identified |
| 79 | 59 | Serum | Mother TN363 | PCV3+ | 208 | none identified |
| 80 | 60 | Fetal lung tissue | Stillborn A from TN363 | PCV3+ | 253 | none identified |
| 81 | 60 | Fetal liver tissue | Stillborn A from TN363 | PCV3+ | 574 | none identified |
| 82 | 61 | Fetal muscle | Stillborn B from TN363 | ND | 824 | none identified |
| 83 | 61 | Fetal heart | Stillborn B from TN363 | PCV3+ | 1425 | none identified |
| 84 | 59 | Placenta | Mother TN363 | PCV3+ | 363 | none identified |
| 85 | 60 | Fetal heart | Stillborn A from TN363 | PCV3+ | 219 | none identified |
